# Transcriptomic comparison of early onset preeclampsia and placenta accreta identifies inverse trophoblast and decidua functions at the maternal-fetal interface

**DOI:** 10.1101/2025.05.09.653201

**Authors:** Ophelia Yin, Ana Almonte-Loya, Romina Appierdo, Monica Yang, Bahar D. Yilmaz, Tomiko T. Oskotsky, Juan M. Gonzalez, Linda C. Giudice, Yalda Afshar, Marina Sirota

## Abstract

Early onset preeclampsia is a placental disorder characterized by shallow implantation, whereas placenta accreta spectrum is a placental disorder of deep placental attachment. This study compares the transcriptome of these two obstetric syndromes. By integrating available microarray and single-cell placenta/decidua transcriptomic datasets, we demonstrated that early onset preeclampsia genes are inversely expressed in placenta accreta, with the most marked differences noted in cell types of decidua, endothelial, and extravillous trophoblasts. Our findings highlight the key functions of trophoblast cell migration and invasion, decidua cell signaling, hypoxia pathways, and global growth factor and collagen contributions to these pregnancy disorders. This research provides new insights into the mechanisms of placentation and unifies these clinical siloes of disease by focusing on the fundamental biology of placental development at the maternal-fetal interface.

## Introduction

Preeclampsia is a hypertensive disorder of pregnancy that affects 5-10% of pregnant people, and it continues to be a significant contributor to both maternal and neonatal mortality worldwide^1^. The origin of preeclampsia remains elusive but is thought to arise from shallow placentation in pregnancy^2^. After embryo implantation, the developing placental cells called trophoblasts attach to the maternal uterine lining (endometrium of pregnancy (“decidua”)). The trophoblasts organize into small finger-like villi projections that interdigitate within the decidua cells and the underlying stromal matrix^3^. Trophoblast cells that extend beyond the villi into the decidua are extravillous trophoblasts, and these play a key role in remodeling the maternal spiral arteries that serve to facilitate nutrient and gas exchange in pregnancy. These extravillous trophoblasts within the vasculature take on an endothelial-like phenotype, migrating in close proximity to the maternal endothelial cells that line the spiral arteries. Abnormalities in the orchestration of placentation are thought to result in increased maternal vascular resistance, decreased oxygen and nutrient exchange, and the resulting endothelial damage and inflammation of preeclampsia that result in systemic manifestations including hypertension, renal damage, liver damage, pulmonary vascular leakage, and cerebral seizures^2,4^. Preeclampsia is defined by these secondary clinical features of a primary placental disorder. Early onset preeclampsia, which occurs prior to 34 weeks of gestation, occurs more rarely but is associated with more severe disease and increased maternal and perinatal morbidity^5,6^.

In stark contrast, placenta accreta spectrum is a placental disorder in which there is such deep villous attachment to the uterus that the placenta does not separate at the time of birth. There is deep remodeling of spiral arteries and neovascularization, leading to the complexity of the condition, requiring preterm delivery via cesarean hysterectomy^7^. Although multiple clinical epidemiologic studies have described opposite clinical manifestations of preeclampsia and placenta accreta^8,9^, these two disorders historically were not believed to share underlying biology. However, we recently conducted a single-cell transcriptomic analysis of placenta accreta and found that many of the genes upregulated in our placenta accreta extravillous trophoblasts, decidua, and endothelial cells are downregulated in preeclampsia studies^10^. This curious molecular parallel was corroborated by Bartels et al.^11^, whose placenta accreta spatial transcriptomic analysis similarly found many significant genes and pathways in accreta that were inversely regulated compared to preeclampsia. However, no prior studies have directly compared the full preeclampsia versus the placenta accreta transcriptome and instead have focused on each disease in isolation with a literature review highlighting a few overlapping genes. Therefore, a comprehensive meta-analysis of both obstetric syndromes is needed to define a reproducible and robust placental signature.

In this study, we sought to compare *in silico* the transcriptomic signatures of early onset preeclampsia and placenta accreta. We targeted early onset preeclampsia due to both its more clinically severe phenotype and its similar preterm birth timing as that of placenta accreta spectrum. The goal of this analysis was to test the hypothesis that these disorders lie on the opposite ends of the spectrum of a shared process of physiologic placentation. We hypothesized that a comparative analysis would narrow down the transcriptional landscape of early onset preeclampsia to a more precise set of genes and pathways relevant to disease etiology or progression^12^. Using publicly available microarrays, we defined an early onset preeclampsia signature and compared to both placenta accreta microarray data and our previous single-cell accreta sequencing data to identify hyper-enriched gene sets and oppositely regulated genes across individual cell types. We found a consistent pattern of enriched and inversely regulated genes when comparing early onset preeclampsia to placenta accreta, identified extravillous trophoblast and decidual genes relevant for cell invasion and antioxidation, and found evidence to support that disorders of placentation are highly dependent on the antecedent uterine environment as a driver of pathologic placentation in pregnancy.

## Results

### Early onset preeclampsia transcriptomic signature and pathway analysis

Using publicly available data, we collated a set of four preeclampsia microarray datasets (GSE25906^13^, GSE74341^14^, GSE75010^15^, GSE93839^16^) across 151 samples meeting our inclusion criteria, see Methods **(Figure 1).** Out of 44 preeclampsia microarrays, 16 were of placental tissue, 10 of these included an appropriate sample size, seven of these included meta-data on mode of delivery, and four of these utilized comprehensive human genome probes to allow for full transcriptomic analysis. We specifically conducted a differential gene expression analysis comparing early onset preeclampsia and preterm controls within these four datasets, as these patients present with earlier and more severe clinical findings with different molecular signatures and maternal morbidity compared with late onset preeclampsia^6,17^. Clinical characteristics of the included patients are detailed in **Table 1**. In total, our study included 86 early onset preeclampsia samples and 65 preterm controls, and we identified 2,600 differentially expressed genes at an adjusted p-value of <0.05, of which 1,239 were upregulated and 1,361 were downregulated for early onset preeclampsia, as shown in the heat map for **Figure 2A**. A volcano plot in **Figure 2B** demonstrates that genes with the highest differential expression (adjusted FDR p-value <0.05, log2FC >0.5) included leptin (*LEP*), high-temperature requirement A-4 (*HTRA4*), follistatin-like 3 (*FSTL3*), type 2 deiodinase (*DIO2*), corticotropin releasing hormone (*CRH*), hexokinase-2 (*HK2*), serpin family A member 3 (*SERPINA3*), transmembrane protein 45A (*TMEM45A*), and SAM and SH3 domain containing 1 (*SASH1*).

**Figure 1:**
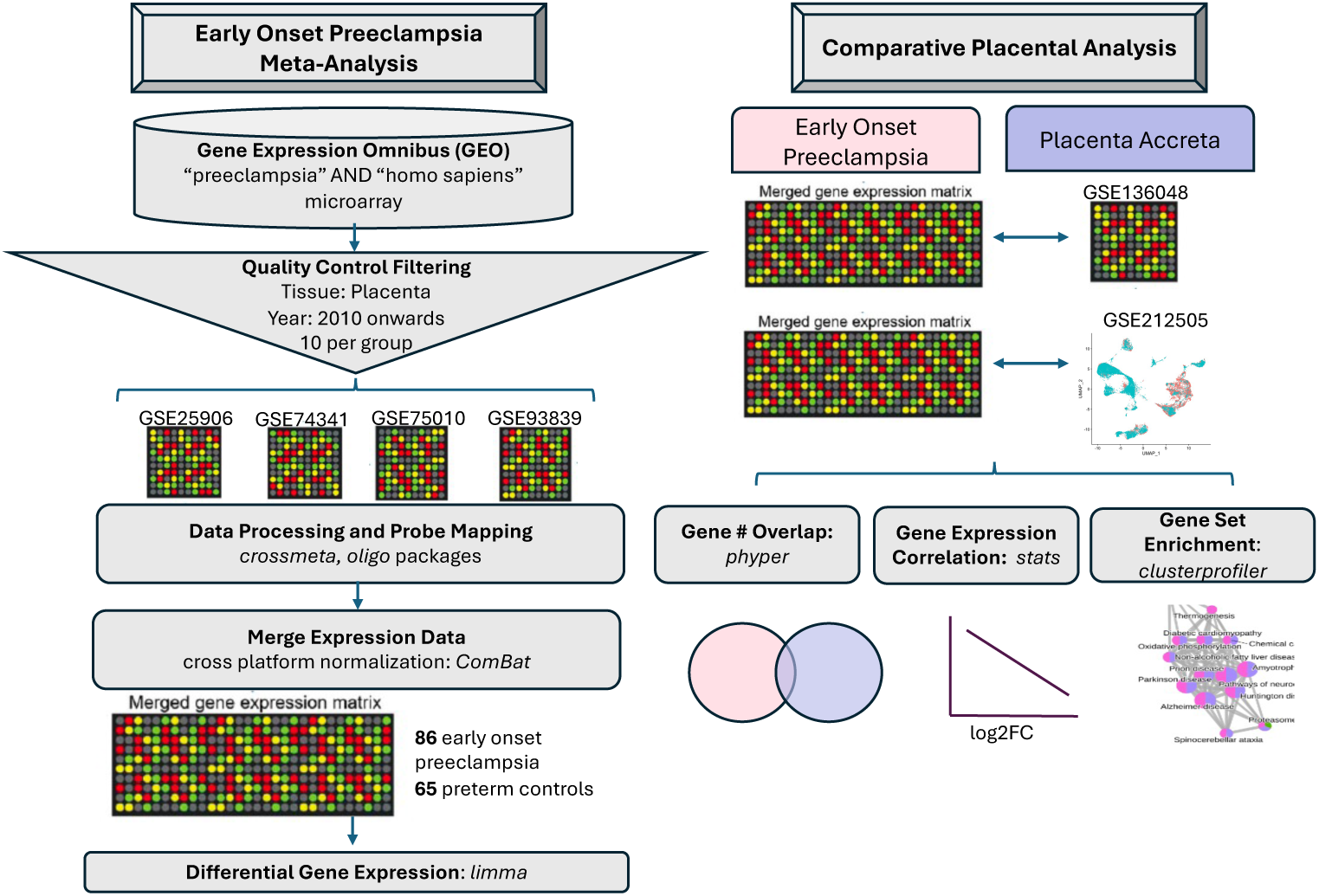
Study methodology overview. Left panel: Schematic representation of methodology for Gene Expression Omnibus microarray dataset search, individual microarray data processing, expression matrix merging, and downstream analyses. Right panel: Schematic representation of the comparative analysis in microarray gene expression between early onset preeclampsia and placenta accreta, comparing the merged preeclampsia expression matrix with that of one preterm accreta microarray (GSE136048) and one placenta accreta single-cell RNA sequencing dataset (GSE212505).

**Figure 2:**
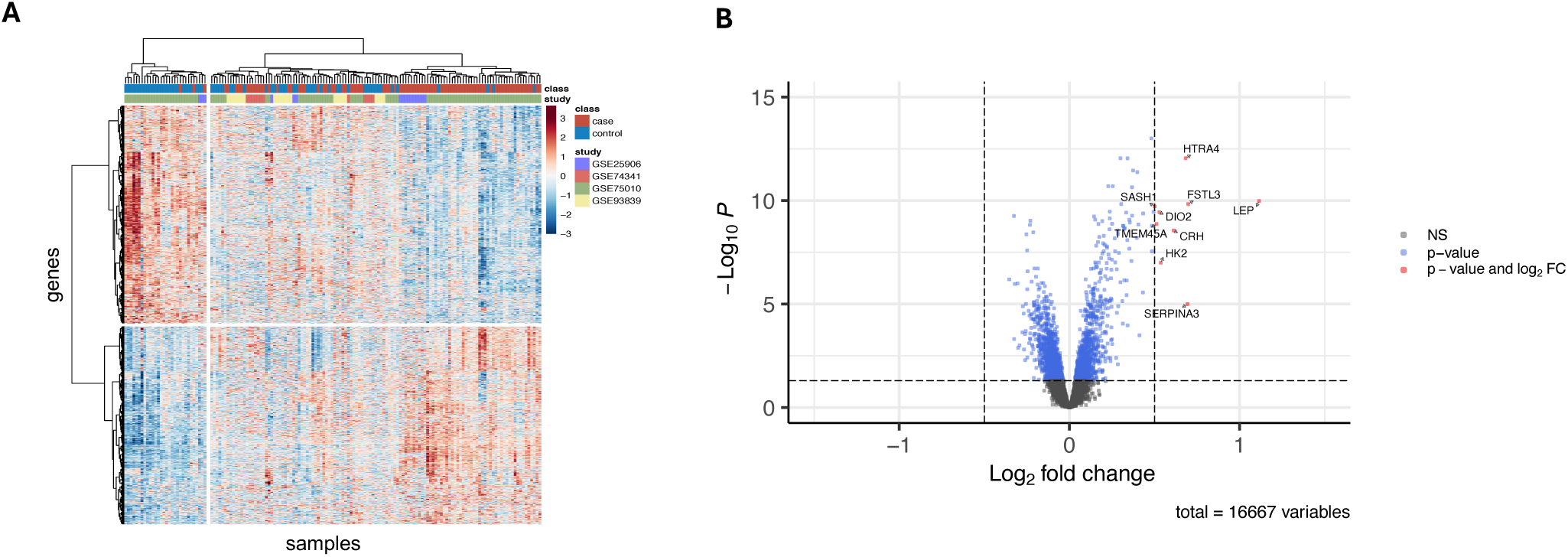
Early onset preeclampsia disease signature. **A** Heat map depicting differential gene expression by case (early onset preeclampsia) and control (preterm control) of the 4 merged microarray studies at an adjusted p-value of <0.05. Analysis demonstrated 2600 differentially expressed genes, of which 1239 are upregulated and 1361 are downregulated. **B** Volcano plot of differential gene expression for early onset preeclampsia, with genes labeled in blue for an adjusted p-value of <0.05 and red for both an adjusted p-value of <0.05 and a log2FC difference of 0.5. NS indicates non-significant.

**Table 1:**
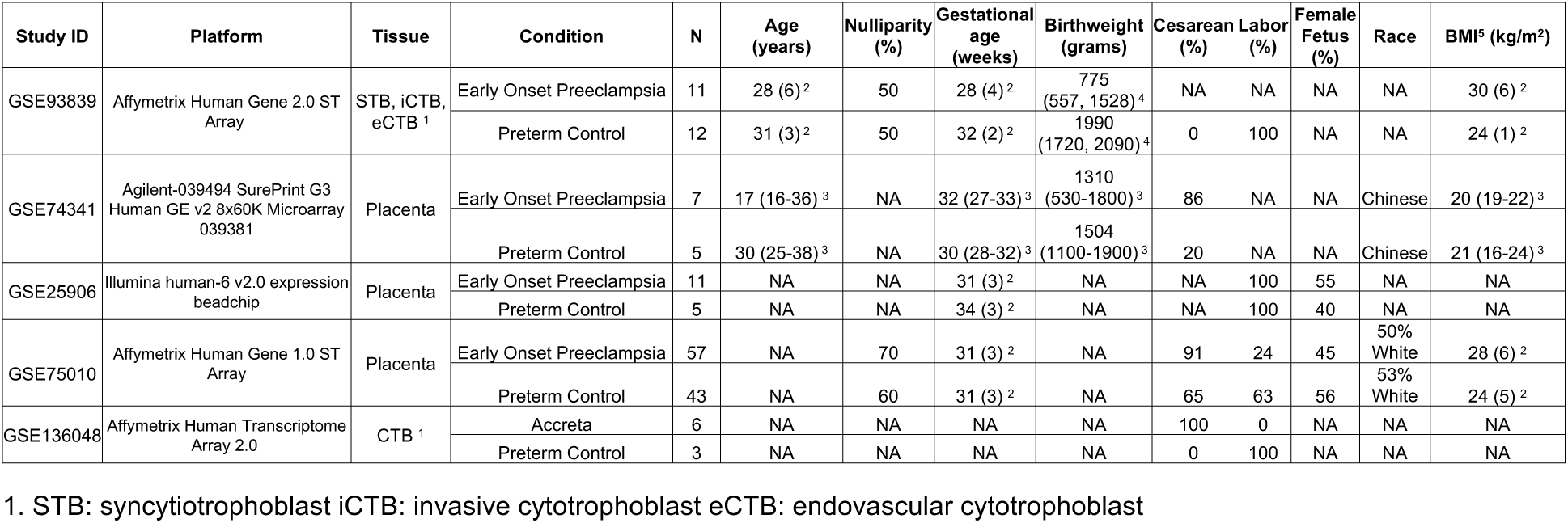
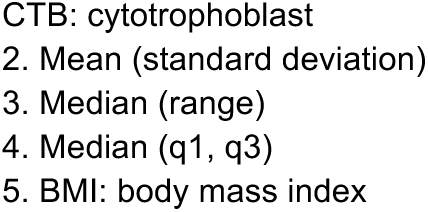
Included microarrays for early onset preeclampsia and placenta accreta with associated metadata.

| Study ID | Platform | Tissue | Condition | N | Age (years) | Nulliparity (%) | Gestational age (weeks) | Birthweight (grams) | Cesarean (%) | Labor (%) | Female Fetus (%) | Race | BMI<sup>5</sup> (kg/m<sup>2</sup>) |
| --- | --- | --- | --- | --- | --- | --- | --- | --- | --- | --- | --- | --- | --- |
| GSE93839 | Affymetrix Human Gene 2.0 ST Array | STB, iCTB, eCTB<sup>1</sup> | Early Onset Preeclampsia | 11 | 28 (6)<sup>2</sup> | 50 | 28 (4)<sup>2</sup> | 775 (557, 1528)<sup>4</sup> | NA | NA | NA | NA | 30 (6)<sup>2</sup> |
|  |  |  | Preterm Control | 12 | 31 (3)<sup>2</sup> | 50 | 32 (2)<sup>2</sup> | 1990 (1720, 2090)<sup>4</sup> | 0 | 100 | NA | NA | 24 (1)<sup>2</sup> |
| GSE74341 | Agilent-039494 SurePrint G3 Human GE v2 8x60K Microarray 039381 | Placenta | Early Onset Preeclampsia | 7 | 17 (16-36)<sup>3</sup> | NA | 32 (27-33)<sup>3</sup> | 1310 (530-1800)<sup>3</sup> | 86 | NA | NA | Chinese | 20 (19-22)<sup>3</sup> |
|  |  |  | Preterm Control | 5 | 30 (25-38)<sup>3</sup> | NA | 30 (28-32)<sup>3</sup> | 1504 (1100-1900)<sup>3</sup> | 20 | NA | NA | Chinese | 21 (16-24)<sup>3</sup> |
| GSE25906 | Illumina human-6 v2.0 expression beadchip | Placenta | Early Onset Preeclampsia | 11 | NA | NA | 31 (3)<sup>2</sup> | NA | NA | 100 | 55 | NA | NA |
|  |  |  | Preterm Control | 5 | NA | NA | 34 (3)<sup>2</sup> | NA | NA | 100 | 40 | NA | NA |
| GSE75010 | Affymetrix Human Gene 1.0 ST Array | Placenta | Early Onset Preeclampsia | 57 | NA | 70 | 31 (3)<sup>2</sup> | NA | 91 | 24 | 45 | 50% White | 28 (6)<sup>2</sup> |
|  |  |  | Preterm Control | 43 | NA | 60 | 31 (3)<sup>2</sup> | NA | 65 | 63 | 56 | 53% White | 24 (5)<sup>2</sup> |
| GSE136048 | Affymetrix Human Transcriptome Array 2.0 | CTB<sup>1</sup> | Accreta | 6 | NA | NA | NA | NA | 100 | 0 | NA | NA | NA |
|  |  |  | Preterm Control | 3 | NA | NA | NA | NA | 0 | 100 | NA | NA | NA |

1. STB: syncytiotrophoblast iCTB: invasive cytotrophoblast eCTB: endovascular cytotrophoblast
CTB: cytotrophoblast 2. Mean (standard deviation) 3. Median (range) 4. Median (q1, q3) 5. BMI: body mass index

Pathway analysis **(Figure 3A,B)** of early onset preeclampsia showed significant activation in biologic processes related to response to hypoxia, regulation of angiogenesis, phosphatidylinositol 3-kinase signaling, and developmental processes involved in reproduction. In contrast, suppression was seen in aerobic and cellular respiration and anabolic pathways of purine ribonucleotide biosynthetic process and sulfur compound metabolic process **(Figure 3A,C)**. An analysis of significantly enriched hallmark pathways additionally found activation of immunologic pathways including tumor necrosis factor-α (TNF-α) signaling, mammalian target of rapamycin complex 1 (mTORC1) signaling, and interleukin2 (IL2) signal transducer and activator of transcription 5 (STAT5) signaling and activation of estrogen response **(Figure 3D,E)**, with suppression of *MYC* proto-oncogene transcription factor targets **(Figure 3D,F)** (all p-adjust <0.05).

**Figure 3:**
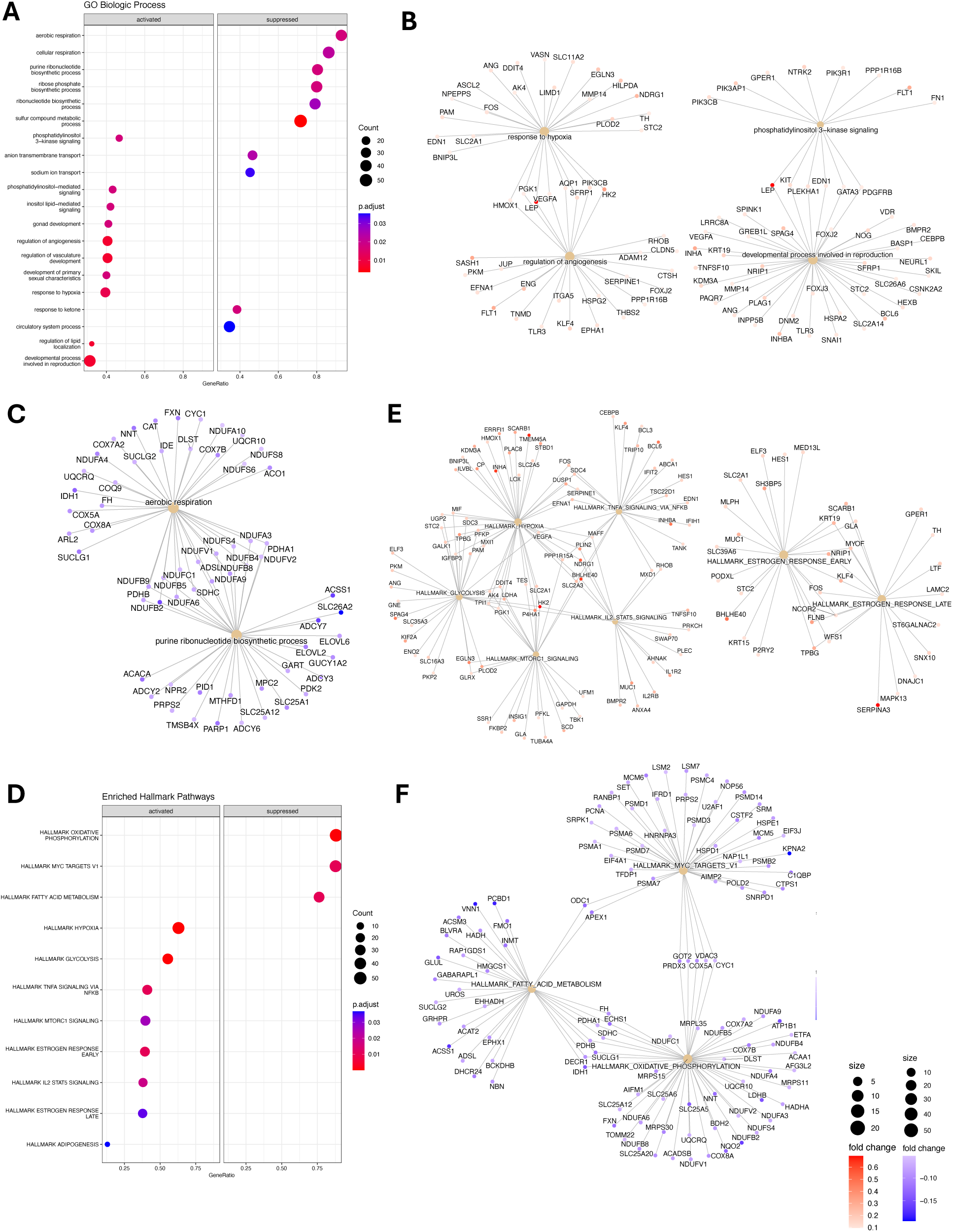
Early onset preeclampsia gene set enrichment analysis. **A** Gene Ontology Biological Process in early onset preeclampsia. **B** Gene-concept network depicting upregulated Gene Ontology Biological Process pathways which are *upregulated*: response to hypoxia, regulation of angiogenesis, developmental process involved in reproduction, phosphatidylinositol 3-kinase signaling. **C** Gene-concept network constructed for the following Gene Ontology Biological Process pathways which are *downregulated*: aerobic respiration and purine ribonucleotide biosynthetic process. **D** Gene set enrichment analysis was performed on Hallmark pathways, with the dotplot indicating significantly enriched pathways in early onset preeclampsia. **E** Gene-concept network constructed for the following Hallmark pathways which are *upregulated:* estrogen response early, estrogen response late, hypoxia, glycolysis, MTORC1 signaling, TNFA signaling via NFKB, IL2 STAT5 Signaling. **F** Gene-concept network constructed for the following Hallmark pathways which are *downregulated:* fatty acid metabolism, oxidative phosphorylation, myc targets.

### Comparative transcriptomic analysis of early onset preeclampsia and placenta accreta Microarray

We next compared our early onset preeclampsia signature with a placenta accreta transcriptome from McNally et al. (GSE136048)^18^, which included 6 accreta placentas and 3 preterm control placentas. To compare the disorders at the same thresholds, we repeated the preeclampsia analysis and the placenta accreta differential gene expression analysis at an unadjusted p-value of <0.05, as described in Methods. For early onset preeclampsia, this resulted in identification of 4,903 differentially expressed genes, of which 2,439 were upregulated and 2,464 were downregulated (**Figure 4A**). For placenta accreta, we identified 416 differentially expressed genes, of which 71 were upregulated and 345 were downregulated **(Figure 4B)**. Comparative analysis of the merged early onset preeclampsia dataset **(Figure 4A)** with the placenta accreta microarray GSE136048 **(Figure 4B)** resulted in 120 shared differentially expressed genes **(Figure 4C)**. The shared genes did not demonstrate hyper-enrichment for early onset preeclampsia genes (p = 0.62). Probing only these shared genes in our early onset preeclampsia dataset and exploring pathway analysis demonstrated that these shared genes are critical in activating catalytic activity and protein metabolic process, while down regulating response to endogenous stimulus, nervous system process, and transmembrane transport (**Figures D, E)**.

**Figure 4:**
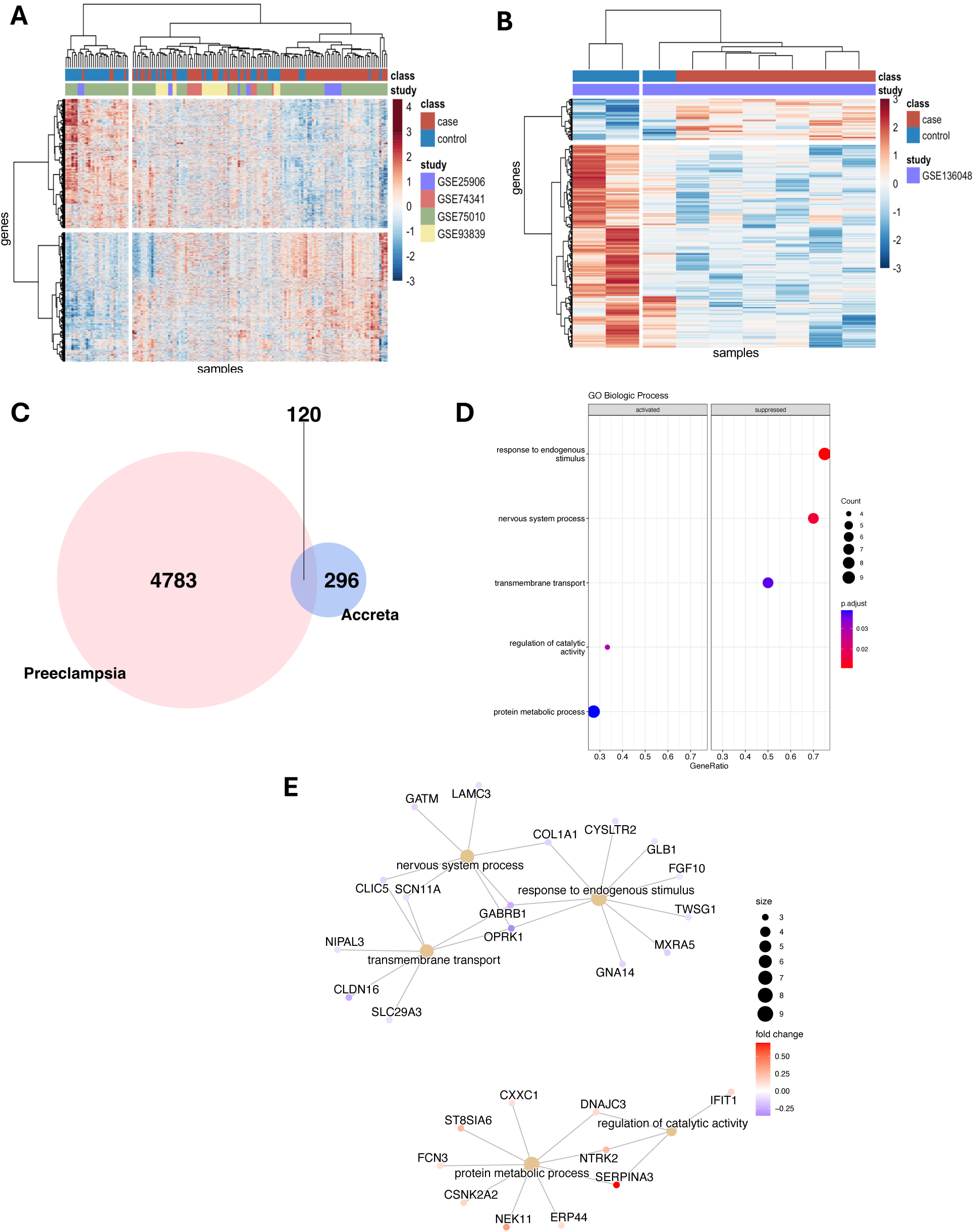
Comparative microarray analysis of early onset preeclampsia and placenta accreta. **A** Early onset preeclampsia: limma analysis demonstrated 4903 differentially expressed genes, of which 2439 are upregulated and 2464 are downregulated. Heat map depicting differential gene expression at an unadjusted p-value of <0.05. **B** Preterm Accreta: limma analysis demonstrated 416 differentially expressed genes, of which 71 are upregulated and 345 are downregulated. Heat map depicting differential gene expression at an unadjusted p-value of <0.05. **C** Venn diagram depicting number of differentially expressed genes that overlap (120 genes) between early onset preeclampsia and preterm accreta, with no hyper enrichment (p=0.62). The total number of genes analyzed was 16,667. **D** Gene Ontology Biological Process in early onset preeclampsia for the 120 overlapping genes. **E** Gene-concept network constructed for the following Gene Ontology Biological Process pathways which are *upregulated*: protein metabolic process, regulation of catalytic activity and *downregulated*: transmembrane transport, nervous system process, response to endogenous stimulus.

When mapping out the log2FC in gene expression among these 120 shared genes, there was a negative correlation in the direction of gene expression (r = -0.39, p < 0.01) **(Figure 5A)**. In addition, when filtering for genes with an absolute difference of at least log2FC of 0.5 between early onset preeclampsia and placenta accreta, we detected nine genes of interest, eight of which demonstrate high expression in early onset preeclampsia but low expression in placenta accreta (ankyrin repeat domain 37 *ANKRD37*, aldehyde oxidase 1 *AOX1*, ceruloplasmin *CP*, guanylate-binding protein 3 *GBP3*, interferon induced protein with tetratricopeptide repeats 1 *IFIT1*, Insulin-like growth factor binding protein 6 *IGFBP6*, NIMA related kinase 11 *NEK11*, and *SERPINA3*) and one with low expression in preeclampsia and high expression in accreta (opioid receptor kappa 1 *OPRK1*) **(Figure 5B).**

**Figure 5:**
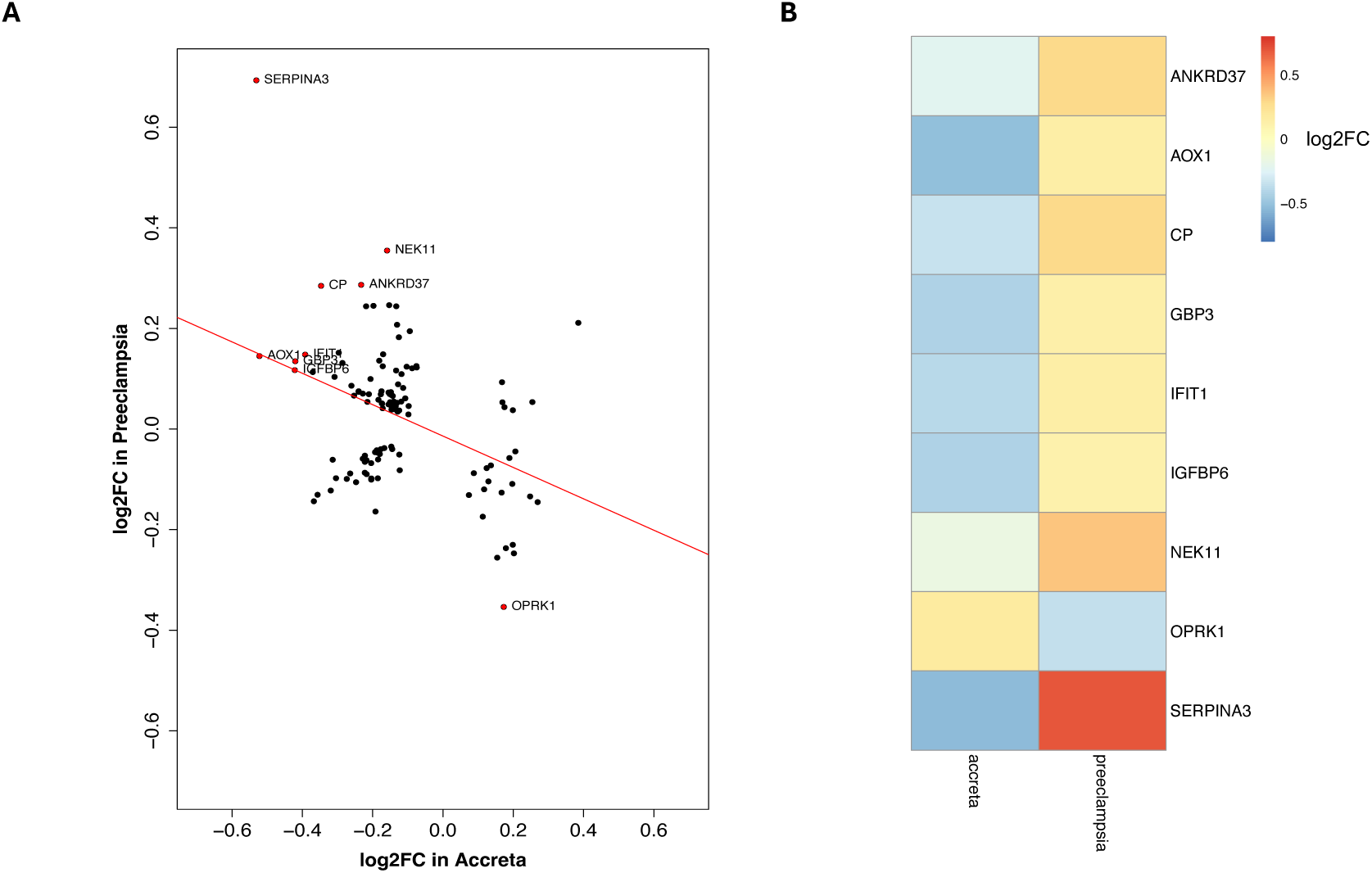
Gene directionality and significant gene comparison for early onset preeclampsia and placenta accreta. A The log2FC in gene expression for early onset preeclampsia is plotted against the log2FC in gene expression for placenta accreta, with *r* = -0.39, p = 8.06e-6. Genes highlighted in red have an absolute difference in log2FC of 0.5 between preeclampsia and accreta. B Heat map comparison of 9 most significantly oppositely regulated genes between early onset preeclampsia and placenta accreta.

### Comparative transcriptomic analysis of early onset preeclampsia and placenta accreta Single-cell

We extended our transcriptomic analysis by comparing our early onset preeclampsia gene signature to a single-cell placenta accreta gene signature, derived from our prior transcriptomic atlas of placenta accreta spectrum (GSE212505)^10^ **(Figure 1)**. In this study, three placenta accreta samples were collected at the time of cesarean hysterectomy at the site of most adherence into the uterine interface and compared to three control placentas collected at the time of cesarean. After isolation of cells from the maternal-fetal interface, the 10x Chromium platform was used to generate libraries for single-cell RNA sequencing and differential gene expression analysis performed using Seurat. In **Table 2**, we show the results of comparing the early onset preeclampsia transcriptomic signature with that of the placenta accreta transcriptome by cell type. Decidua type 1, decidua type 3, endothelial cells, and extravillous trophoblasts from placenta accreta demonstrate hyper enrichment and negative correlation of shared differentially expressed genes with early onset preeclampsia. We previously identified decidua type 1 as perivascular decidua cells, mapping close to endothelial cells **(Supplementary Figure 5)**, while decidua type 3 are decidua cells at the decidua-myometrial interface^10^. The consistent pattern of negative correlation in gene expression when comparing early onset preeclampsia and placenta accreta shared genes, by cell type, is visualized in **Figure 6**.

**Figure 6:**
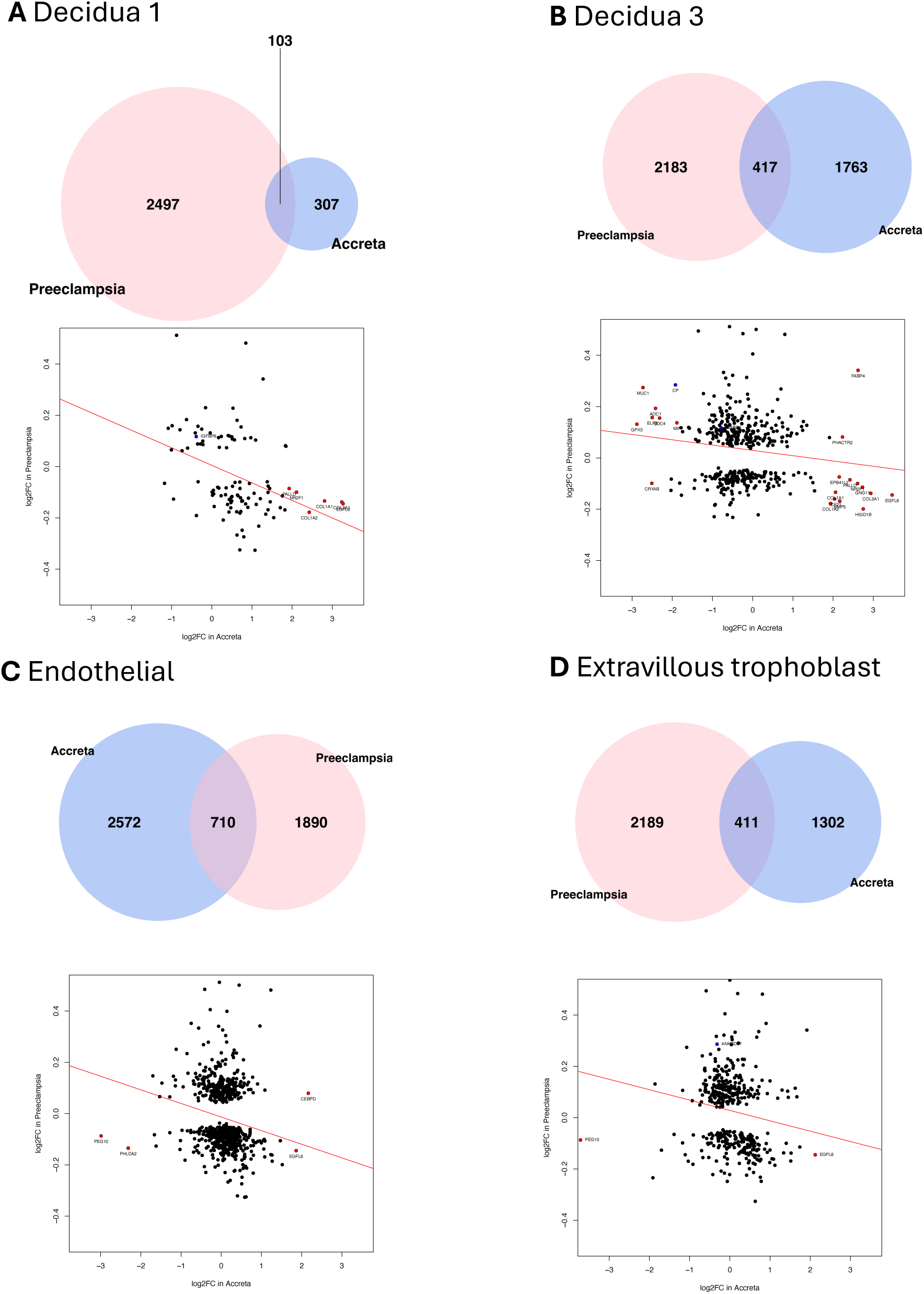
Comparative transcriptomic analysis of early onset preeclampsia and placenta accreta, by cell type. A *Decidua 1:* Overlap of 103 genes. Significant genes with a log2FC of >2 between preeclampsia and accreta include *COL1A1, COL1A2, COL3A1, EGFL6, NR2F1, PALLD* (red)*. IGFBP6* is graphed and represents the overlap between the significant microarray data and this single-cell analysis (blue). B *Decidua 3:* Overlap of 417 genes. Significant genes with a log2FC of >2 between preeclampsia and accreta include *AOC1, BMP5, COL1A1, COL1A2, COL3A1, CP, CRYAB, EGFL6, ELF3, EPB41L2, FABP4, GNG11, GPX3, HIGD1B, MIF, MUC1, NR2F1, PALLD, PHACTR2, SDC4*, and *TCF21* (red). *CP* and *IGFBP6* is graphed and represents the overlap between the significant microarray data and this single-cell analysis (blue). C *Endothelial:* Overlap of 710 genes. Significant genes with a log2FC of >2 between preeclampsia and accreta include *CEBPD, EGFL6, PEG10,* and *PHLDA2* (red). No overlap between the significant microarray data and this single-cell analysis (blue). D *Extravillous trophoblast:* Overlap of 411 genes. Significant genes with a log2FC of >2 between preeclampsia and accreta include *EGFL6* and *PEG10* (red). *ANKRD37* is graphed and represents the overlap between the significant microarray data and this single-cell analysis (blue).

**Table 2:** Comparative analysis of early onset preeclampsia merged dataset compared to single-cell placenta accreta differential gene expression, by cell type.

| Placenta Cell Type | # Accreta Differentially Expressed Genes | # Overlap Genes with Early Onset Preeclampsia | Hyper enrichment p-value | Correlation coefficient *r* | Correlation p-value |
| --- | --- | --- | --- | --- | --- |
| **Decidua 1** | **410** | **103** | **3.04 x 10<sup>-7\*</sup>** | **-0.38** | **7.11 x 10<sup>-5\*</sup>** |
| Decidua 2 | 25 | 10 | 0.003* | -0.49 | 0.150 |
| **Decidua 3** | **2180** | **417** | **1.12 x 10<sup>-6\*</sup>** | **-0.13** | **8.00 x 10<sup>-3\*</sup>** |
| **Endothelial** | **3282** | **710** | **5.58 x 10<sup>-25\*</sup>** | **-0.18** | **9.20 x 10<sup>-7\*</sup>** |
| Cytotrophoblast | 51 | 20 | 3.88 x 10<sup>-5*</sup> | -0.42 | 0.060 |
| Syncytiotrophoblast | 1343 | 278 | 1.28 x 10<sup>-7*</sup> | -0.01 | 0.840 |
| **Extravillous trophoblast** | **1713** | **411** | **6.17 x 10<sup>-22\*</sup>** | **-0.16** | **0.002\*** |
| Lymphoid | 31 | 13 | 4.00 x 10<sup>-4*</sup> | -0.42 | 0.160 |
| Myeloid | 222 | 50 | 4.00 x 10<sup>-4*</sup> | -0.15 | 0.290 |

\*p < 0.05. Cell type rows with both significant gene set hyper enrichment and negative correlation are bolded.

Of the microarray shared genes from **Figure 5B**, *IGFBP6* in decidua type 1 and decidua type 3 cells **(Figure 6 A, B)**, *CP* in decidua type 3 cells (**Figure 6B)**, and *ANKRD37* in extravillous trophoblasts **(Figure 6D)** demonstrated higher expression in early onset preeclampsia bulk tissue than placenta accreta single-cell analysis.

Other genes of interest with a log2FC difference of at least 2 between early onset preeclampsia and placenta accreta included a number of collagen genes which were downregulated in preeclampsia (*COL1A1*, *COL1A2, COL3A1*) compared to placenta accreta decidua cells **(Figure 6A)**, epidermal growth factor-like domain multiple-6 (*EGFL6*) which was downregulated in preeclampsia compared to all accreta cell types **(Figure 6)**, and paternally expressed gene 10 (*PEG10*) which was relatively upregulated in preeclampsia compared to accreta endothelial and extravillous trophoblast cells (**Figure 6C, D)**. Overall, decidua type 3 had the highest number of genes that show highly disparate and negatively correlated expression values compared to early onset preeclampsia.

Pathway analysis of the shared genes between early onset preeclampsia and decidua type 3 **(Figure 7A)** and endothelial cells **(Figure 7B)** demonstrate overlapping pathways between preeclampsia and accreta that may govern normal decidual signaling and endothelial migration at the maternal-fetal interface. These include the hallmark pathways of hypoxia, glycolysis, and early estrogen response. We also noted downregulation of oxidative phosphorylation that may be due to exposure of decidua cells to a low oxygen environment in the setting of reduced placental attachment or decidual resistance.

**Figure 7:**
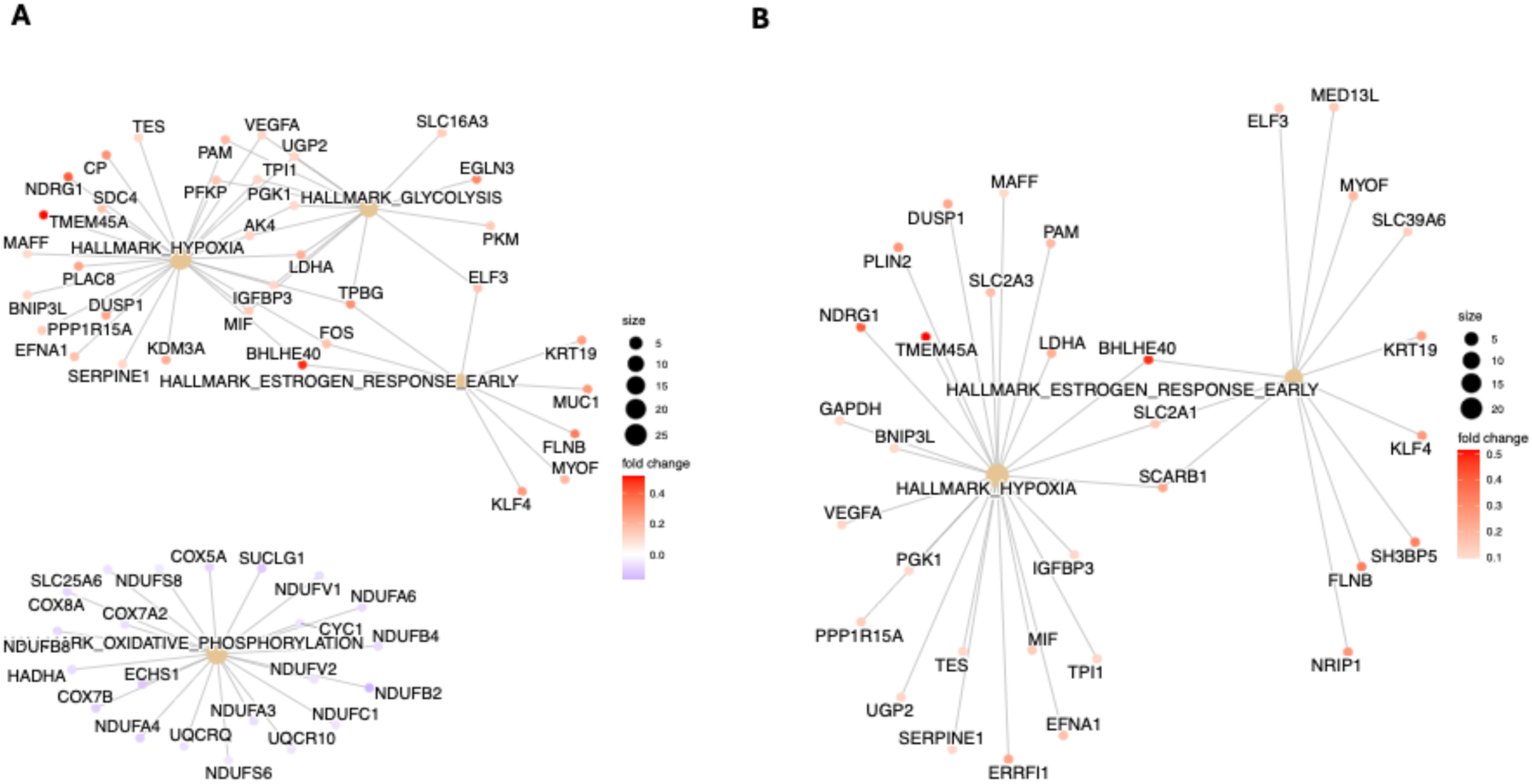
Pathway analysis of overlapping genes between early onset preeclampsia and placenta accreta. A Decidua 3: Gene-concept network constructed for the Hallmark pathways B Endothelial: Gene-concept network constructed for the Hallmark pathways *Other cell types of interest (decidua 1, extravillous trophoblast) did not demonstrate gene set enrichment*.

## Discussion

Early onset preeclampsia and placenta accreta, despite their distinct clinical manifestations, may represent opposite extremes of the physiologic placentation process. To explore this hypothesis, we leveraged bulk and single-cell placental transcriptomic signatures and performed a comparative analysis of early onset preeclampsia with placenta accreta to identify the key gene sets and pathways for these conditions. We computationally merged four early onset preeclampsia datasets, using their respective meta-data to understand and account for clinical covariates. Our differential gene expression analysis of 86 early onset preeclampsia and 65 preterm control placentas resulted in identification of multiple genes and pathways of interest. Candidate genes including *LEP, HTRA4, DIO2,* and *CRH* were significantly upregulated in our study and these same genes were also identified in a separate study by Ma et al.^19^, who compared preterm preeclampsia to preterm controls using independent datasets GSE44711 and GSE66273. Their study validated *ANKRD37*, *CRH*, and *LEP* as diagnostic markers with AUC values of >0.9 when tested in a third dataset (GSE4707) and confirmed both mRNA and protein expression of these genes in separate human placental samples. The reproducibility of our findings in an independent study using distinct and smaller datasets highlights their robustness, broad biological relevance, and their ability to translate into actionable diagnostic tools in clinical practice.

Extending our comprehensive early onset preeclampsia meta-analysis, we then proceeded with a comparative analysis of early onset preeclampsia with placenta accreta and propose several candidate genes for further exploration that may regulate the spectrum of normal placentation, given that they were both shared in these placental pathologies and oppositely regulated.

### Extravillous trophoblast genes of interest include ANKRD37, SERPINA3, and PEG10

The prevailing history of preeclampsia has been a study of trophoblasts, as the placenta is often the easiest specimen to obtain in clinical research. In our study, we identified three trophoblast genes of interest: *ANKRD37*^20^ and *SERPINA3*^21^ which were upregulated and *PEG10* which was downregulated in early onset preeclampsia. *In vitro* studies suggest that ANKRD37 plays a crucial role in the hypoxia response and may contribute to the development of preeclampsia under hypoxic conditions^22,23^. Moreover, studies of *ANKRD37* in extravillous trophoblast cells^20^ demonstrated that it prevents cell migration and invasion by suppressing matrix metalloproteases, upregulating adhesion molecules such as E-cadherin, and possibly signaling via the NF-κB pathway^20^. Similar upregulation of E-cadherin has been seen in neighboring syncytiotrophoblasts^24^. In our single-cell analysis, *ANKRD37* localized to extravillous trophoblasts with minimal expression in placenta accreta but relatively high expression in early onset preeclampsia, highlighting that this gene may be involved in minimizing the degree of placental attachment to the maternal decidua. *SERPINA3* demonstrated the highest preeclampsia expression compared to accreta in our microarray analysis, but while it has been shown to be upregulated in placentas complicated by preeclampsia^21,25^, its function is paradoxically to promote cell invasion. Since *SERPINA3* is induced under hypoxic conditions, its upregulation may be a marker of low oxygen conditions in the late trimester preeclampsia placenta as opposed to serving as a cause of disease pathogenesis. *PEG10* expression is decreased in early onset preeclampsia, with negative regulation in settings of hypoxia and TNF-α inflammation, but it has not been shown to affect cell differentiation or proliferation^26^. Taken together, these genes represent interesting candidates for functional perturbation in both preeclampsia and accreta models.

### Decidual genes of interest include IGFBP6, BMP5, MUC1, SDC4, and NR2F1

While much research on preeclampsia has focused on trophoblast populations, emerging data emphasize the importance of decidual resistance in both predisposing to and resulting in a pregnancy with preeclampsia^27–29^. Conversely, in the setting of accreta, there is iatrogenic injury, which leads to absent or defective decidua and therefore is predominantly seen in patients with a prior cesarean delivery and placenta previa^30^. In our study, the greatest number of oppositely and differentially regulated genes between early onset preeclampsia and placenta accreta were in the decidual cell populations, adding more evidence that it may be the environment of implantation that drives the clinical sequelae of placental disorders. For example, we found that *IGFBP6* was upregulated in early onset preeclampsia and downregulated in accreta decidual cells. A recent single-cell study of decidual tissue collected from patients with a history of early onset preeclampsia also defined *IGFBP6* as a marker of early onset preeclampsia^27^. Interestingly, multiple genes identified in our decidual population have been shown to play a role in cell proliferation, invasion, migration, or angiogenesis when studied in trophoblasts, but their function in the decidua are less clear. These include bone morphogenic protein 5 (*BMP5*)^31^, mucin 1 (*MUC1*)^32^, syndecan-4 (*SDC4*)^33^, and nuclear receptor subfamily 2 group F member 1 (*NR2F1*)^34^. Instead of studying specific genes in extravillous trophoblast models in isolation, these data suggest that decidua-trophoblast co-culture, placental explants in decidual cell environments, or organoid/assembloid models^35^ may better recapitulate the biologic niche of the maternal-fetal interface and uncover insights into both preeclampsia and accreta pathology.

### The role of hypoxia in decidua signaling and CP function

Hypoxia pathways and their downstream effects may also be important in trophoblast attachment to the decidua. Previous studies demonstrated a lack of Hypoxia-inducible factor 1-alpha (HIF-1α) expression in decidua cells from invasive placental accreta compared to control, implicating that hypoxia signaling serves to stem placental growth during implantation. We identified multiple activated hypoxia pathways from early-onset preeclampsia placentas and found that *CP* is upregulated in preeclampsia and downregulated in placenta accreta specifically in the decidual cell population. *CP* is a copper-containing ferroxidase that reduces oxidative damage under low oxygen conditions. *CP* levels are increased in placentas of severe preeclampsia, localizing to the intervillous space and syncytiotrophoblasts, and CP increases in blood with worsening clinical features of preeclampsia^36–38^. Further study of *CP*’s role and the general role of hypoxia in decidual signaling may help define pathways that regulate placentation early in development.

### Global decidual contribution of EGFL6 and Collagen in the Extracellular Matrix

We observed a global decrease in *EGFL6* in early onset preeclampsia, with concomitant increase in gene expression for placenta accreta across decidua, endothelial, and extravillous trophoblast cell types. A recent single-cell preeclampsia study similarly noted downregulation of *EGFL6* in stromal cells from early onset preeclampsia^31^, though the role of this gene has not been functionally explored. Finally, we found notable downregulation of collagen (*COL1A1*, *COL1A2*, and *COL3A1*) in early onset preeclampsia and upregulation in placenta accreta, a finding that was also found in single-cell analysis of stromal cells from early onset preeclampsia^39^. Previous placenta accreta studies have noted a lack of *TGF-*β staining at sites of excessive decidual and myometrial collagen, indicating that changes in the extracellular matrix may serve to dampen boundary signals and facilitate cell invasion.

### Summary

In summary, our comparative analysis of early onset preeclampsia with placenta accreta identified trophoblast genes responsible for cell migration and invasion and highlighted the global contribution of *EGFL6* and collagen at the maternal-fetal interface. Critically, we defined the most overlapping and inversely regulated genes among the decidua cell populations when comparing preeclampsia to accreta, emphasizing the importance of the environment of implantation as a driver of clinical sequelae in placental disorders. A schematic of our findings is summarized in **Figure 8**.

**Figure 8:**
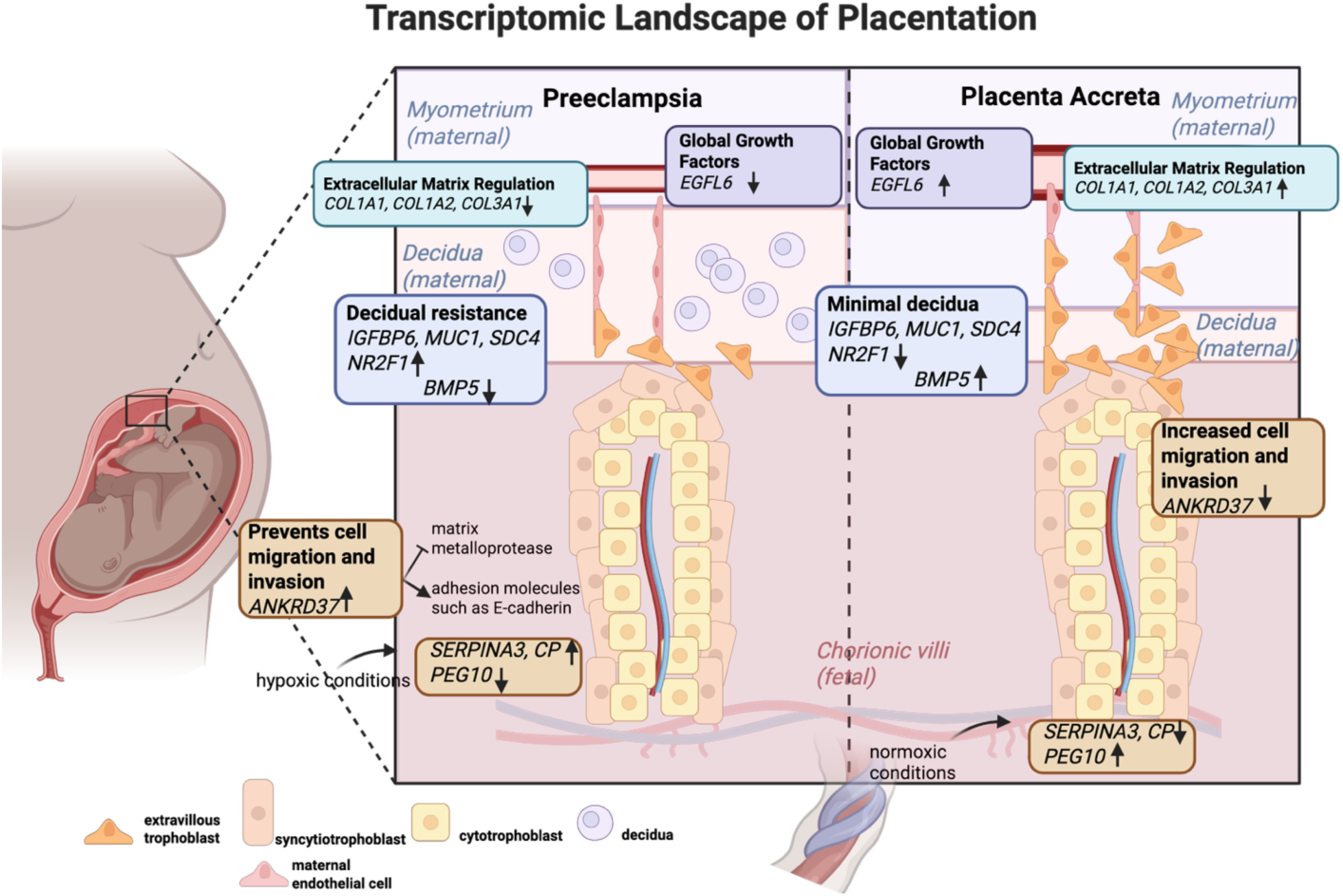
Summary figure. Created in https://BioRender.com

While our study provides valuable insights into the opposing mechanisms in early onset preeclampsia and placenta accreta, several limitations must be acknowledged. There were a small number of placenta accreta datasets available for comparison (one microarray and one single-cell dataset), and the small number of clinical samples in each of these data (9 and 6, respectively). The study focuses primarily on early onset preeclampsia and placenta accreta, which may not capture the full spectrum of these conditions, particularly late onset preeclampsia or cases of placenta accreta spectrum disorders with varying degrees of severity. Our analysis assumes that the etiology of both conditions is primarily related to the placenta and immediately attached decidua cells, without the ability to address potential maternal or systemic factors that may also contribute to the pathophysiology^40^. This assumption simplifies the complex interplay between maternal and fetal contributions to these disorders and leaves more to be explored in relation to the decidua and deeper uterine layers. The reliance on transcriptomic data, although robust, does not account for post-transcriptional modifications, protein-level changes, or functional validation of the identified pathways, which limits the direct clinical applicability of these findings. Additionally, while we computationally merged datasets to improve statistical power and reduce batch effects, the heterogeneity of patient populations, clinical presentations, and sample collection protocols across datasets may introduce variability that is difficult to fully control.

### Future directions

We should aim to move away from categorizing placental disorders into discrete clinical silos and instead adopt flexibility in seeing placentation, from embryo implantation and miscarriage to placental growth and accreta as a spectrum of obstetric disease to bring more resources and scientists together in defining the fundamental biology of placentation. Future work can extend this analysis by exploring not only the genes discussed, but also those genes identified with no prior data that may be new targets for diagnostic and therapeutic discovery. We plan to explore additional cell-specific comparisons using emerging single-cell datasets^27,39^ with additional emphasis on the decidua environment in pathologic placentation.

## Methods

An overview of the biocomputational workflow is detailed in **Figure 1**. In brief, we utilized publicly available preeclampsia microarrays to create a merged dataset to define differentially expressed genes in early onset preeclampsia compared to preterm controls. We then probed this signature with a placenta accreta microarray dataset and then our previous single-cell placenta accreta data to understand shared genes and pathways between early onset preeclampsia and placenta accreta. We tested for gene set enrichment, gene expression correlation, and utilized gene set enrichment analysis to generate gene-concept maps of the most relevant biologic and hallmark pathways.

### Data collection and dataset merging

The NCBI Gene Expression Omnibus (GEO) (http://www.ncbi.nlm.nih.gov/geo/) was searched for all publicly available preeclampsia datasets using the keywords “preeclampsia” AND “homo sapiens”, searching for data types “expression profiling by array, by high throughput sequencing, by sage” by November of 2023. We included microarrays from 2010 onwards of placental tissue collected at the time of delivery. We excluded datasets with less than 10 samples per group. Additionally, datasets were excluded if they were poorly annotated or utilized array platforms with limited number of probes. The search yielded four preeclampsia datasets available for analysis. Similar search was used to query placenta accreta datasets, of which only one microarray dataset was available. Clinical covariates of age, parity, gestational age, cesarean birth, labor, fetal sex, race, and body mass index (BMI) were collected, as available.

Raw data from each microarray dataset was downloaded and processed using R version 4.2.3. and the Bioconductor version 3.16 with packages *crossmeta*, *oligo*^41^, *biomatr*^42,43^, and *limma*^44^. Each microarray underwent processing including background correction, log2-transformation, intra-study quantile normalization, and probe gene mapping from respective probe IDs to gene symbols. All Affymetrix, Agilent, and Illumina platforms were processed using *crossmeta* except GSE75010, which required processing using the *oligo* package due to its historical array platform (Human Gene 1.0 ST Array).

Merging and cross-platform normalization of the five datasets (four preeclampsia and one accreta) was carried out using ComBat within the R package *sva*^45^. The principal component analysis plots before and after normalization are shown in **Supplementary Figure 1**. After merging, the total number of overlapping genes was 16,667 (**Supplementary Figure 2**).

### Clinical covariate selection

We conducted a principal component analysis for each clinical covariate of interest to assess its importance in modifying global gene expression in our merged dataset. We utilized Levene’s test for statistical analysis of equality of variances. The results are shown in **Supplementary Figure 3**. Based on these findings, we selected cell type as a covariate in our differential gene expression analysis. While mode of delivery showed significant variance, we further examined whether laboring status prior to birth was different in our cohorts, regardless of whether they achieved a cesarean or vaginal birth. Since there was no difference in expression by laboring status, we decided not to adjust for mode of delivery. Prior studies have also described the lack of significant gene perturbation based on labor status^46^.

We excluded gestational age from the covariate analysis as previous analyses have demonstrated a strong influence of trimester on gene expression in the placenta^47,48^. To avoid this confounding effect, we decided to include only early onset preeclampsia patients (delivering at gestational age ≤ 34 weeks) and corresponding preterm control patients (delivering at ≤ 37 weeks) for our merged microarray analysis. Including preterm patients also allowed for a better comparison with the placenta accreta dataset, as accreta patients are recommended to deliver at ≤ 34 weeks gestation to prevent maternal morbidity^49^. Studies describing indication for preterm birth in their control group noted that the delivery was spontaneous and for non-infectious causes^16,18^.

### Early onset preeclampsia differential gene expression and pathway analysis

We conducted a differential gene expression analysis using *limma* on the subset of the merged dataset containing the four preeclampsia microarrays to define the disease signature of early onset preeclampsia. Cell type was used as a covariate in the linear model using an adjusted false discover rate (FDR) p-value of < 0.05. *pheatmap* and *EnhancedVolcano* were used to generate plots of significant genes, with the volcano plot utilizing a log fold change (log FC) cut-off of 0.5. Gene set enrichment analysis was conducted using *clusterProfiler*^50^ and *DOSE*^51^. Visualization of enriched terms and gene-concept networks were created using *enrichplot*.

### Comparative gene analysis of early onset preeclampsia and placenta accreta Microarray transcriptome

Differential gene expression analysis was repeated on the placenta accreta microarray (GSE136048). An unadjusted p-value of <0.05 was used to account for the small number of samples, as analyzed in the original published study^18^. Differential gene expression on the four preeclampsia microarrays was also performed at an unadjusted p-value of <0.05 to allow for comparative gene analysis. We quantified the number of overlapping genes and performed pathway analysis on the shared genes. To assess statistical significance of gene enrichment between early onset preeclampsia and placenta accreta, we used hypergeometric testing accounting for the 16,667 background genes. Correlation between log2FC in gene expression between preeclampsia and placenta accreta was assessed using Pearson’s correlation testing.

### Single-cell transcriptome

Our placenta accreta single-cell dataset (GSE212505)^10^ was previously processed by applying quality filters and filtering doublets. It was also normalized, scaled and integrated using Harmony to remove potential batch effects. Previous clusters were found with FindNeighbors and FindClusters (resolution = 0.5). We re-analyzed this previously processed single-cell placenta accreta dataset (GSE212505)^10^ to include only the comparison between placenta accreta-adherent samples and control samples, excluding the internal control of placenta accreta-non adherent samples. We adjusted our analysis based on sample batch and sample timing of delivery (term vs preterm), as not all controls were preterm in this study (**Supplementary Figure 4).** Analysis was performed with *Seurat* using the FindMarkers function with MAST as the testing method, incorporating batch and time of delivery as latent variables in the differential expression model^52^. We utilized the same cell markers for cell annotation as in our prior analysis and performed differential gene expression for the placenta accreta disease signature by cell type, with an adjusted FDR p-value of <0.05. **(Supplementary Figure 5).** The number of cells and differentially expressed genes are listed in **Supplementary Table 1.** We then repeated our comparative analysis, comparing each gene signature by cell type for placenta accreta with the merged early onset preeclampsia disease signature with an adjusted FDR p-value of <0.05.

## Supporting information

Meta-analysis differential gene expression preterm preeclampsia

Differential gene expression analysis placenta accreta GSE136048

Single cell accreta differential gene expression decidua 1

Single cell accreta differential gene expression decidua 3

Single cell accreta differential gene expression extravillous trophoblast

Single cell accreta differential gene expression endothelial

120 overlapping preeclampsia and accreta genes

Preeclampsia and single cell accreta decidua 1 overlap

Preeclampsia and single cell accreta decidua 3 overlap

Preeclampsia and single cell accreta extravillous trophoblast overlap

Preeclampsia and single cell accreta endothelial overlap

## Data Availability

The datasets used in this study are publicly available from the NCBI Gene Expression Omnibus (GEO) (http://www.ncbi.nlm.nih.gov/geo/). All data is accessible under public access terms. The code used for this study, including pre-processing, analysis and visualization, can be accessed via our GitHub repository: https://github.com/oyin128/Transcriptomic-comparison-of-early-onset-preeclampsia-and-placenta-accreta.

## Acknowledgements

This work was funded by the National Institutes of Health (NIH) Eunice Kennedy Shriver National Institute of Child Health and Human Development (NICHD) [P01HD106414, R01HD105256] and the March of Dimes Prematurity Research Center at UCSF [60982053-50185]. Ophelia Yin was supported in part by the Chan Zuckerberg Biohub Physician-Scientist Fellowship Program, as part of the Chan Zuckerberg Initiative. The funders played no role in the study design, data collection, analysis and interpretation of data, or the writing of this manuscript.

## Author contributions

O.Y., A.A.L., R.A. contributed to the study design, analysis, and table and figure creation. M.Y., B.D.Y., T.T.O, J.M.G., L.C.G., Y.A., and M.S. contributed to the study design and study interpretation. O.Y., A.A.L., R.A., T.T.O, and M.S. wrote the main manuscript text. All authors reviewed the manuscript.

