## Supplementary Information for "Transcriptomic comparison of early onset preeclampsia and placenta accreta identifies inverse trophoblast and decidua functions at the maternal-fetal interface"

### Supplementary Figure 1: Harmonization of five microarray datasets

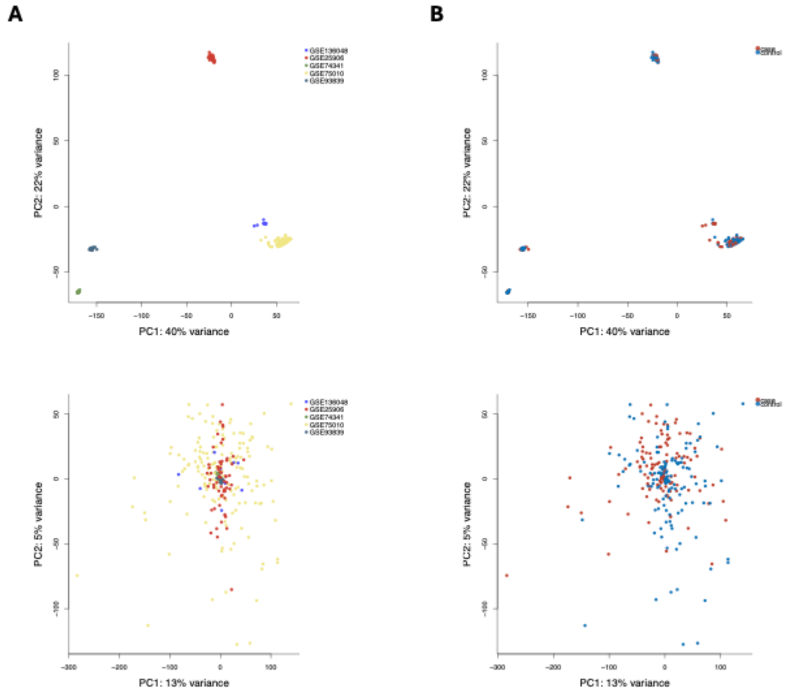


A Merging, with study labeled
B Merging, with case and control labeled

### Supplementary Figure 2: Gene overlap in merged dataset

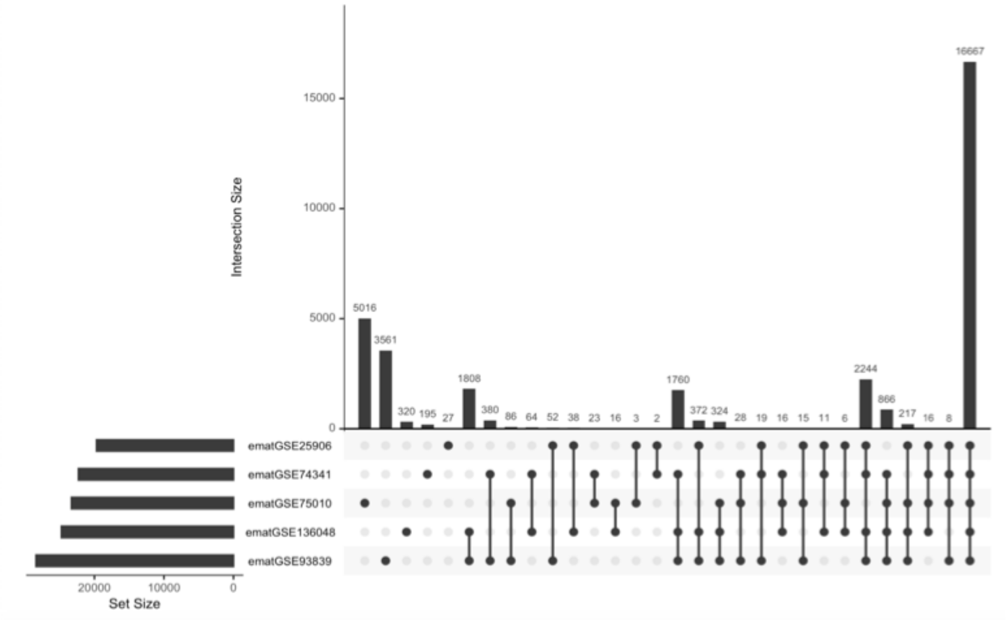


Upset plot demonstrating the 5 microarray datasets and the overlap in gene set size for each one, resulting in a total of 16,667 shared genes across the 5 microarrays.

### Supplementary Figure 3: PCA plots to explore variance of covariates in merged dataset

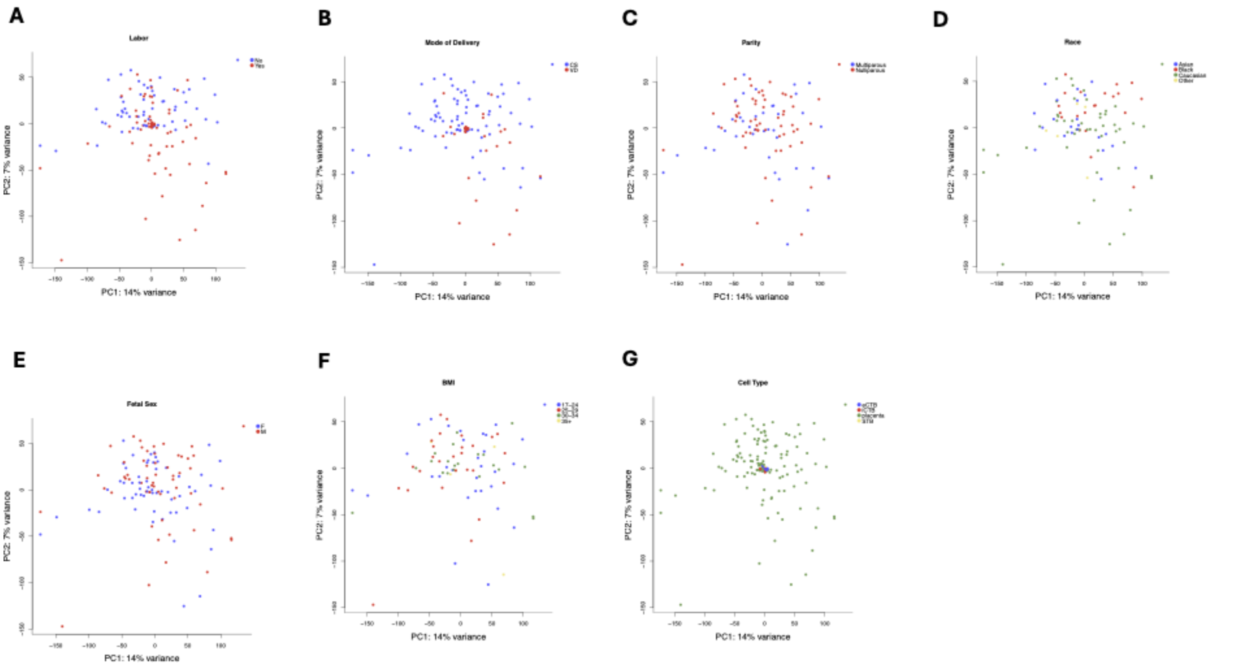


A Labor, p>0.05. B Mode of delivery, p=0.02. CS: cesarean section. VD: vaginal delivery. C Parity, p>0.05. D Race, p>0.05. E Fetal sex, p>0.05. F: female. M: male. F Body mass index (BMI, in kg/m$^2$), p>0.05.
G Cell type, p<0.01. eCTB: endovascular cytotrophoblast. iCTB: invasive cytotrophoblast. STB: syncytiotrophoblast.

### Supplementary Figure 4: Covariate correction for placenta accreta single-cell data.

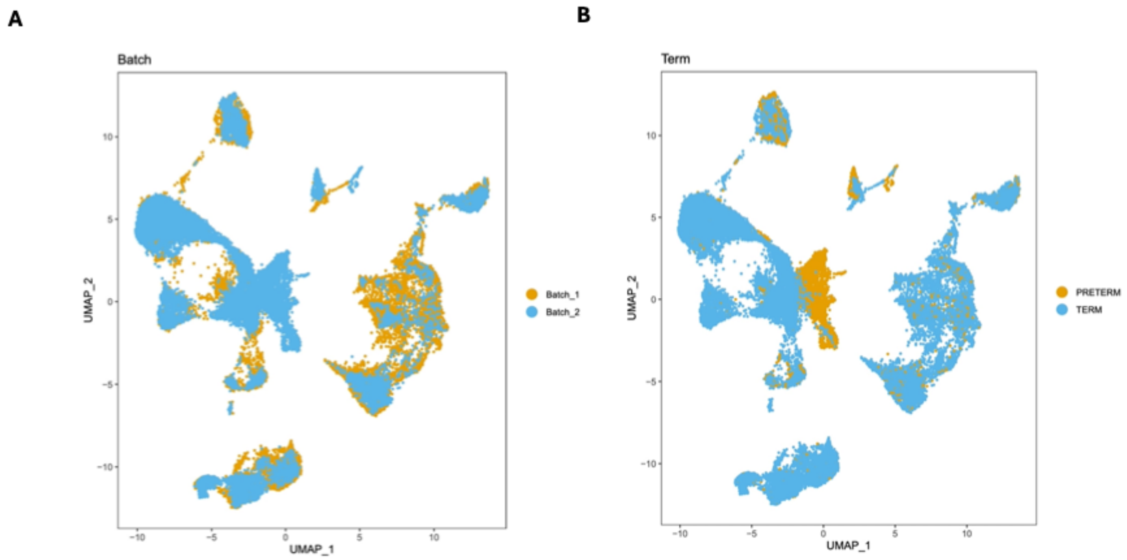


Placenta accreta single-cell UMAP data with mapping of individual cells by A batch and B preterm or term. Both covariates were adjusted for in the final differential gene expression analysis for the single-cell data.

### Supplementary Figure 5: Results of single-cell placenta accreta analysis.

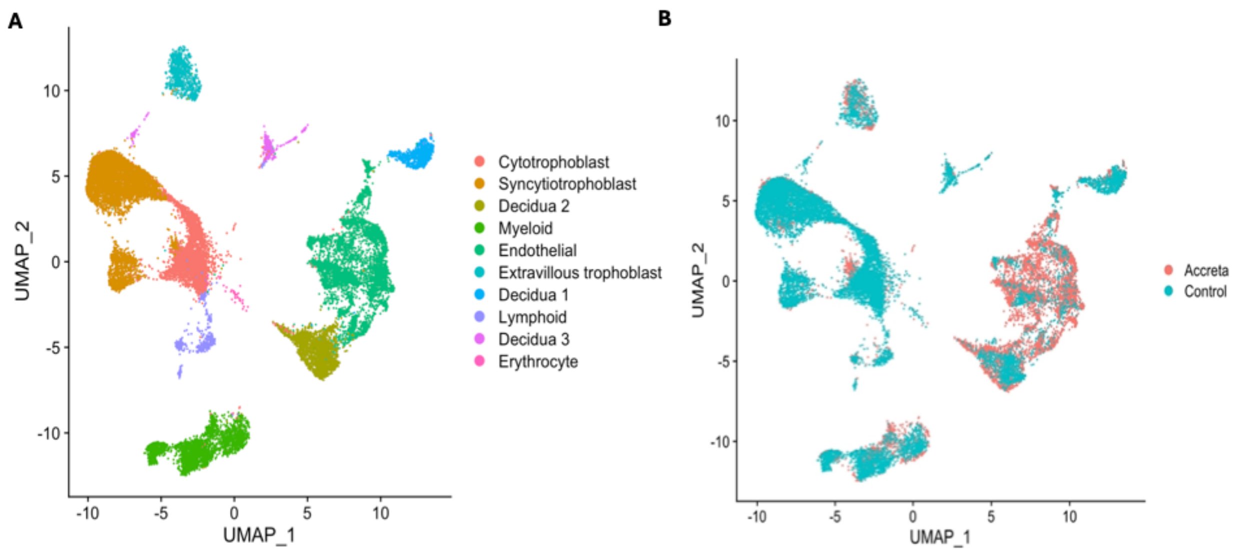


A UMAP identifying cell type designation associated with gene clusters.
B UMAP representation split by disease phenotype of placenta accreta (n=3) and control (n=3).

### Supplementary Table 1: Single-cell accreta analysis.

| Cell Type | Cell Number | Up # | Down # | Total Gene # |
|---|---|---|---|---|
| Cytotrophoblast | 6531 | 32 | 19 | 51 |
| Decidua1 | 1326 | 265 | 145 | 410 |
| Decidua2 | 2435 | 20 | 5 | 25 |
| Decidua3 | 913 | 782 | 1398 | 2180 |
| Endothelial | 6158 | 2023 | 1259 | 3282 |
| Erythrocyte | 602 | 0 | 0 | 0 |
| Extravillous trophoblast | 1637 | 780 | 933 | 1713 |
| Lymphoid | 1048 | 7 | 24 | 31 |
| Myeloid | 3963 | 137 | 85 | 222 |
| Syncytiotrophoblast | 6793 | 627 | 716 | 1343 |

Number of cells and differentially expressed genes identified by cell based on disease phenotype for placenta accreta, indicating positive or negative expression compared to control.

**Supplementary Data Excel Sheets**

**metalimmapreeclampsiacontrol_preterm_preEonly_all.csv**

Meta-analysis differential gene expression results for 4 preterm preeclampsia datasets (GSE25906, GSE74341, GSE75010, GSE93839) adjusted for cell type

**metalimmaGSE136048.csv**

Differential gene expression result for 1 placenta accreta dataset GSE136048

**celltype_DEG.csv x 10**

Placenta accreta single cell differential gene expression results- for each cell type, adjusted for batch and term.

**metalimmaoverlap_2_120.csv**

120 genes shared between preterm preeclampsia and placenta accreta arrays.

**celltype_8.2.24.csv x 10**

Genes shared between preterm preeclampsia and single-cell placenta accreta analysis, by cell type.
